# Structural basis for backtracking by the SARS-CoV-2 replication-transcription complex

**DOI:** 10.1101/2021.03.13.435256

**Authors:** Brandon Malone, James Chen, Qi Wang, Eliza Llewellyn, Young Joo Choi, Paul Dominic B. Olinares, Xinyun Cao, Carolina Hernandez, Edward T. Eng, Brian T. Chait, David E. Shaw, Robert Landick, Seth A. Darst, Elizabeth A. Campbell

## Abstract

Backtracking, the reverse motion of the transcriptase enzyme on the nucleic acid template, is a universal regulatory feature of transcription in cellular organisms but its role in viruses is not established. Here we present evidence that backtracking extends into the viral realm, where backtracking by the SARS-CoV-2 RNA-dependent RNA polymerase (RdRp) may aid viral transcription and replication. Structures of SARS-CoV-2 RdRp bound to the essential nsp13 helicase and RNA suggested the helicase facilitates backtracking. We use cryo-electron microscopy, RNA-protein crosslinking, and unbiased molecular dynamics simulations to characterize SARS-CoV-2 RdRp backtracking. The results establish that the single-stranded 3’-segment of the product-RNA generated by backtracking extrudes through the RdRp NTP-entry tunnel, that a mismatched nucleotide at the product-RNA 3’-end frays and enters the NTP-entry tunnel to initiate backtracking, and that nsp13 stimulates RdRp backtracking. Backtracking may aid proofreading, a crucial process for SARS-CoV-2 resistance against antivirals.

**Significance Statement:** The COVID-19 pandemic is caused by the severe acute respiratory syndrome coronavirus 2 (SARS-CoV-2). The SARS-CoV-2 genome is replicated and transcribed by its RNA-dependent RNA polymerase (RdRp), which is the target for antivirals such as remdesivir. We use a combination of approaches to show that backtracking (backwards motion of the RdRp on the template RNA) is a feature of SARS-CoV-2 replication/transcription. Backtracking may play a critical role in proofreading, a crucial process for SARS-CoV-2 resistance against many antivirals.

## Introduction

SARS-CoV-2 is the causative agent of the current COVID-19 pandemic (1, 2). The SARS-CoV-2 genome is replicated and transcribed by its RNA-dependent RNA polymerase holoenzyme [holo-RdRp, subunit composition nsp7/nsp8_2_/nsp12 (3, 4)] in a replication-transcription complex (RTC), which is the target for antivirals such as remdesivir [Rdv; (5)]. The holo-RdRp is thought to coordinate with many co-factors to carry out its function (6, 7). Some of these co-factors, such as the nsp13 helicase (8) and the nsp10/nsp14 proofreading assembly (9, 10), are also essential for viral replication and are antiviral targets (11–13).

We recently reported the first views of the SARS-CoV-2 RTC in complex with the nsp13 helicase [cryo-electron microscopy structures at a nominal resolution of 3.5 Å; (14)]. The overall architecture of the nsp13-RTC places the nucleic acid binding site of nsp13 directly in the path of the downstream template-strand RNA (t-RNA), and cryo-electron microscopy (cryo-EM) difference maps revealed the 5’-single-stranded t-RNA overhang engaged with nsp13 before entering the RdRp active site (14). The nsp13 helicase translocates on single-stranded nucleic acid in the 5’->3’ direction (15–22). Thus, this structural arrangement presents a conundrum: The RdRp translocates in the 3’->5’ direction on the t-RNA strand, while nsp13 translocates on the same strand in the opposite direction. Translocation of each enzyme opposes each other, and if the helicase prevails it is expected to push the RdRp backward on the t-RNA (14). This reversible backward sliding, termed backtracking, is a well-studied feature of the cellular DNA-dependent RNA polymerases [DdRps; (23–30)].

Backtracking by the cellular DdRps plays important roles in transcription regulation, including the control of DdRp pausing during transcription elongation, termination, DNA repair, and transcription fidelity (25). In backtracking, the DdRp and associated transcription bubble move backwards on the DNA, while the RNA transcript reverse threads through the complex to maintain the register of the RNA-DNA hybrid (23–30). This movement generates a single-stranded 3’- segment of the RNA transcript which is extruded out the secondary or NTP-entry tunnel that branches off from the primary DdRp active-site cleft around the conserved bridge helix (27–31).

Although evolutionarily unrelated to the DdRps, a secondary channel, formed by the RdRp motif F β-hairpin loop and proposed to serve as an NTP-entry tunnel, branches off from the main SARS-CoV-2 RdRp active site channel (32). This NTP-entry tunnel is well positioned to receive the single-stranded 3’- segment of backtracked RNA, a structural architecture analogous to the DdRps (14). We envisaged that translocation by the helicase could mediate backtracking of the RdRp, an otherwise energetically unfavorable process, enabling the key viral functions such as proofreading (9, 10, 12, 33) and template switching during subgenomic transcription (7, 34). Here we outline the structural basis for SARS-CoV-2 RTC backtracking and describe the role of nsp13 in stimulating backtracking.

## Results

### SARS-CoV-2 RdRp backtracked complexes for cryo-electron microscopy

Previously, DdRp backtracked complexes (BTCs) were generated for structural studies by direct incubation of the DdRp with DNA-RNA scaffolds containing mismatched nucleotides at the RNA 3’-end (27, 28, 30); these BTC-scaffolds bind with the downstream Watson-Crick base pairs of the RNA-DNA hybrid positioned in the DdRp active site and the single-stranded 3’-segment of mismatched RNA extruding out the DdRp NTP-entry tunnel. To study RdRp BTCs, we therefore designed and tested RNA scaffolds based on our original SARS-CoV-2 RTC-scaffold but with three or five mismatched cytosine nucleotides added to the product-RNA (p-RNA) 3’-end (BTC_3_- and BTC_5_- scaffolds; Fig. 1A).

**Fig. 1.**
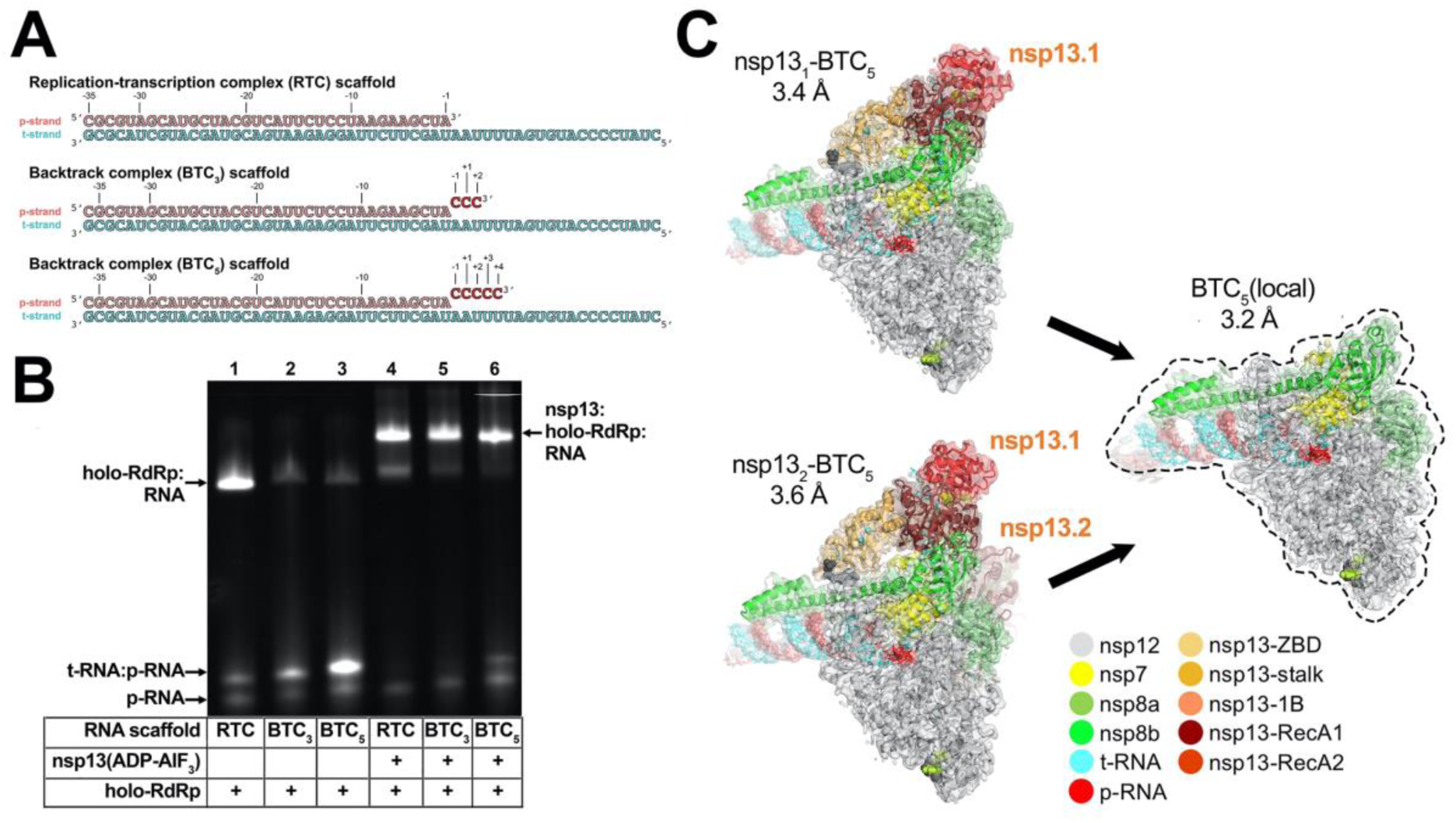
SARS-CoV-2 backtrack complex. **A.** RNA scaffolds: (*top*) replication-transcription complex (RTC) scaffold (14); (*bottom*) backtrack complex scaffolds (BTC_3_ and BTC_5_). **B.** A native gel electrophoretic mobility shift assay reveals that holo-RdRp requires nsp13(ADP-AlF_3_) to bind the BTC scaffolds efficiently. **C.** Cryo-EM structures of SARS-CoV-2 BTCs. Shown is the transparent cryo-EM density [local-resolution filtered; (47)] with the refined models superimposed (Table S1). The models and density are colored according to the key.

Native electrophoretic mobility shift assays revealed that although the holo-RdRp (nsp7/nsp8_2_/nsp12) bound the RTC-scaffold as observed previously [Fig. 1B, lane 1; SI Appendix, Fig. S1A; (14)], nsp13 was required for efficient binding to the BTC-scaffolds (Fig. 1B). Stable nsp13-holo-RdRp complexes with BTC- scaffolds were also observed by native mass-spectrometry (Fig. S1B and C).

To determine the structural organization of the SARS-CoV-2 BTC, we assembled nsp13(ADP-AlF_3_) and holo-RdRp with the BTC_5_-scaffold (Fig. 1A; hereafter called BTC_5_) and analyzed the samples by single-particle cryo-EM. The sample comprised two major classes: nsp13_1_-BTC_5_ (3.4 Å nominal resolution), and nsp13_2_-BTC_5_ (3.6 Å; Fig. 1C; SI Appendix, Fig. S2 and S3). To eliminate structural heterogeneity in the nsp13 subunits and obtain a higher-resolution view of the BTC, the particles from both classes were combined and locally refined inside a mask applied around the holo-RdRp and RNA (excluding the nsp13 subunits), leading to the BTC_5_(local) combined map (3.2 Å; Fig. 1C; SI Appendix, Fig. S2 and S3, Table S1).

The cryo-EM maps (Fig. 1C and 2) revealed two significant differences with the nsp13-RTC structures (14): 1) The single-stranded downstream template-RNA (t-RNA) engaged with nsp13.1 was resolved (Fig. 2A), and 2) a single-stranded p-RNA 3’-segment was extruded into the RdRp NTP-entry tunnel (Fig. 2B).

**Fig. 2.**
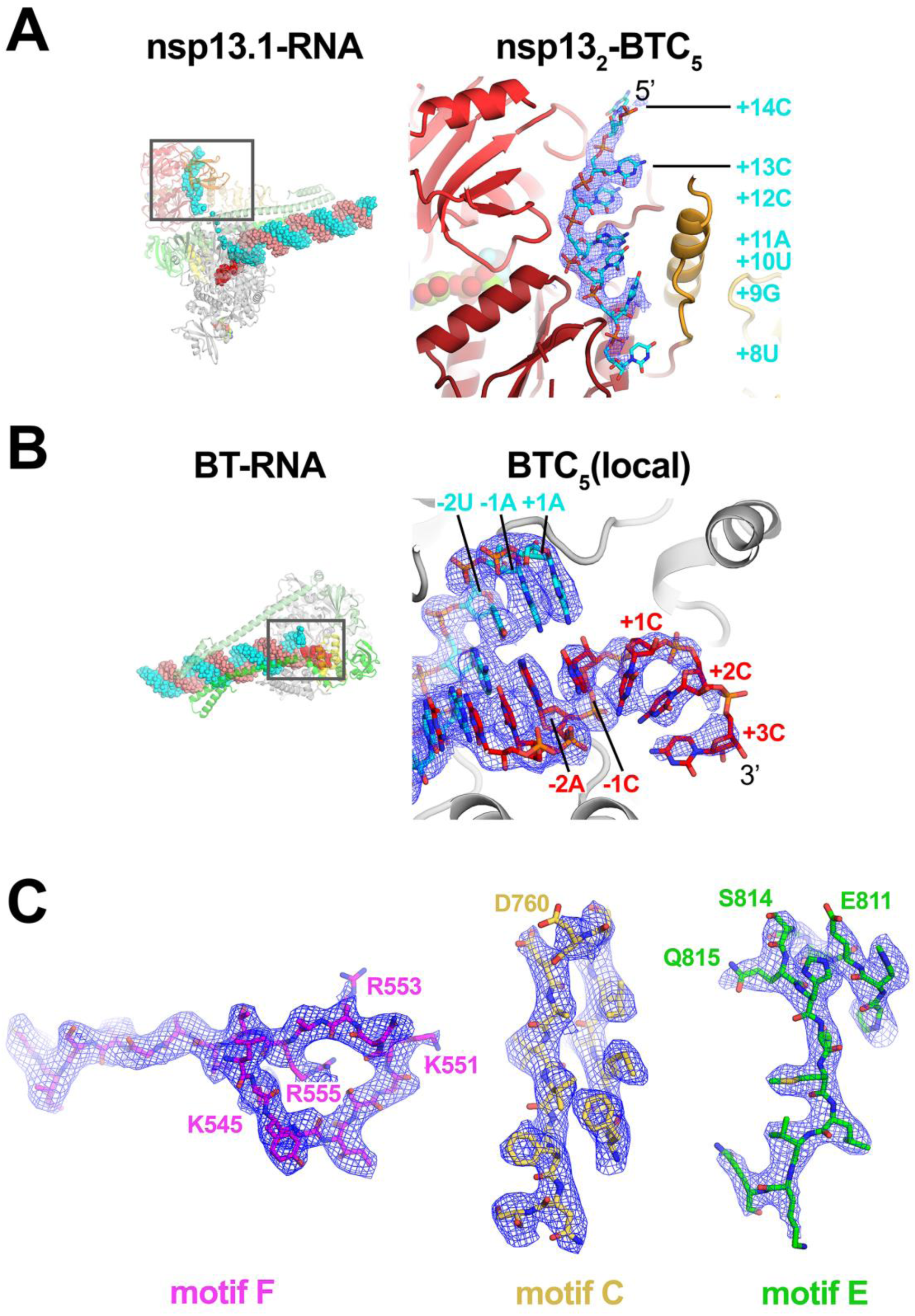
Cryo-EM density maps. **A.** (*left*) Overall view of nsp13_1_-BTC_5_. The boxed region is magnified on the right. (*right*) Magnified view of the t-RNA segment (+14-5’-CCCAUGU-3’-+8) enclosed in the nsp13.1 helicase subunit. The cryo-EM density map (from the nsp13_2_-BTC structure) for the RNA is shown (blue mesh). **B.** (*left*) Overall view of the BTC structure. The boxed region is magnified on the right. (*right*) Magnified view of the region around the RdRp active site, showing the t-RNA (cyan) and p-RNA (red) with the backtracked RNA segment. The cryo-EM density map for the RNA [from BTC_5_(local)] is shown (blue mesh). **C.** BTC_5_(local) cryo-EM density maps around nsp12 conserved motifs F, C, and E. Selected residues are labeled.

### Nsp13 binds the downstream single-stranded t-RNA

In the nsp13_1_-BTC_5_ and nsp13_2_-BTC_5_ cryo-EM maps, the single-stranded 5’-segment of the t-RNA was engaged with nsp13.1. This region of the cryo-EM density was well-resolved (Fig. 2A), allowing identification of the t-RNA segment engaged within the helicase as +14 to +8 (numbering defined in Figure 1A), ^5’^CCCAUGU^3’^. The five-nucleotide segment connecting the t-RNA between the helicase and the RdRp (+7 to +3) was disordered and not modeled.

### The SARS-CoV-2 RdRp NTP-entry tunnel accommodates the backtracked RNA

The cryo-EM maps also resolved a single-stranded p-RNA 3’-segment of the BTC_5_-scaffold extruding into the RdRp NTP-entry tunnel (Fig. 2B), confirming the formation of a BTC (Fig. 3A). The overall architecture of the SARS-CoV-2 BTC is analogous to DdRp BTCs [Fig. 3; (14)]. The DdRp bridge helix [BH; (35)] separates the DdRp active site cleft into a channel for the downstream template DNA (over the top of the BH; Fig. 3B) and the NTP-entry tunnel (underneath the BH; Fig. 3B). Similarly, the viral RdRp motif F [SI Appendix, Fig. S4A; (32)] serves as the strand separating structural element for the backtracked RNA (Fig. 3A). The downstream t-RNA passes over the top of motif F, while the backtracked RNA extrudes out the NTP-entry tunnel underneath motif F (Fig. 3A).

**Fig. 3.**
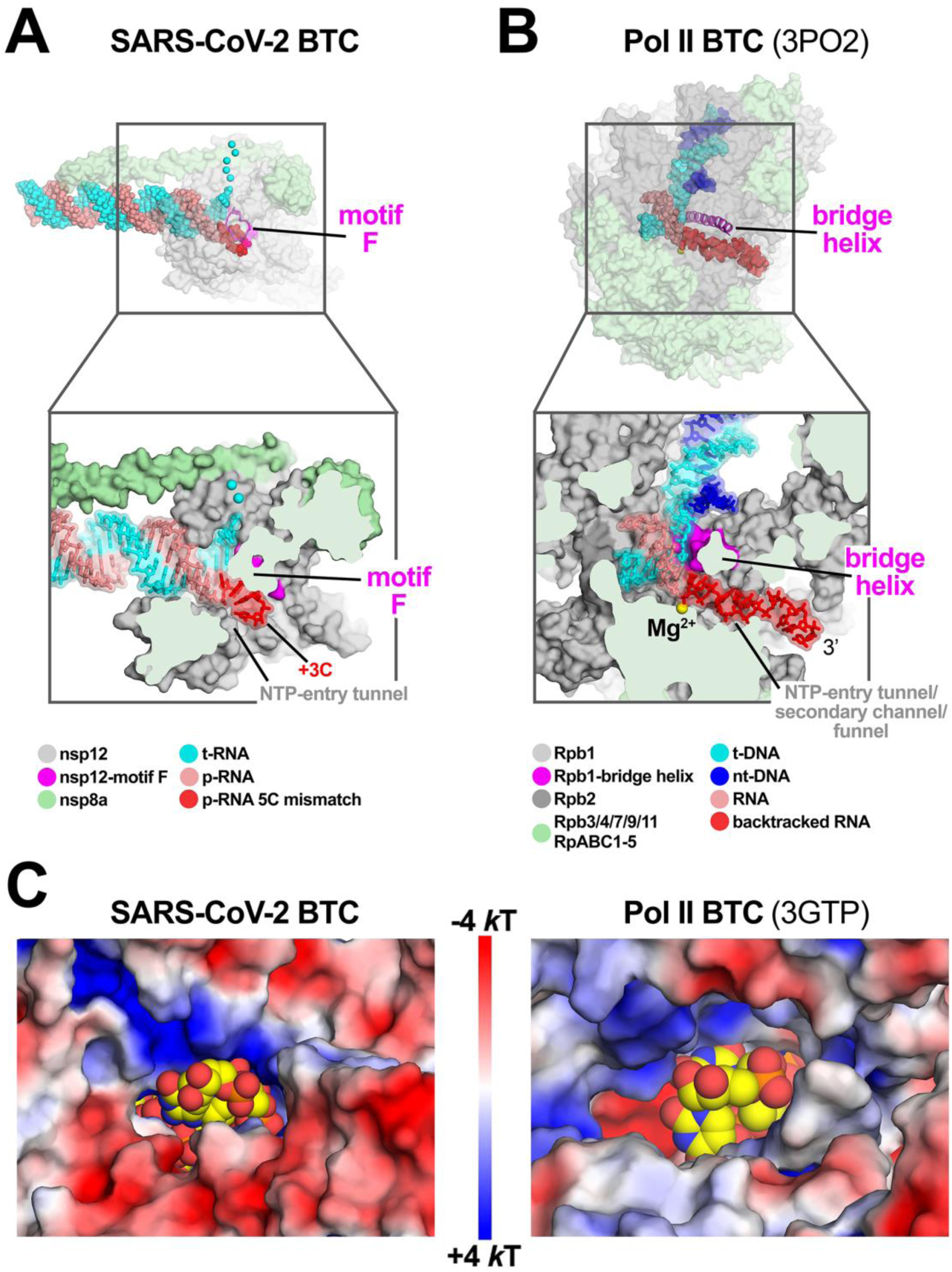
SARS-CoV-2 RdRp and DdRp BTCs. **A, B.** SARS-CoV-2 RdRp (**A**) and DdRp (**B**) BTCs. (*top*) Proteins are shown as transparent molecular surfaces, nucleic acids as atomic spheres. The boxed regions are magnified on the bottom. (*bottom*) Magnified, cross-sectional view. Proteins are shown as molecular surfaces, nucleic acids in stick format with transparent molecular surface. **A.** The SARS-CoV-2 BTC_5_(local). Nsp8a and nsp12 are shown (nsp7 and nsp8b are removed for clarity). Nsp12 motif F is shown as a magenta backbone ribbon (*top*). Backtracked RNA (+1C to +3C of the BTC_5_-scaffold; Figure 1A) extrude out the NTP-entry tunnel. **B.** A DdRp (*Saccharomyces cerevisiae* Pol II) BTC [PDB ID: 3PO2; (29)]. The bridge helix is shown as a magenta backbone ribbon. The backtracked RNA extrudes out the NTP-entry tunnel/secondary channel/funnel. **C.** Views from the outside into the NTP-entry tunnels of the SARS-CoV-2 (*left*) and an *S. cerevisiae* DdRp [PDB ID: 3GTP; (27) BTC. Protein surfaces are colored by the electrostatic surface potential [calculated using APBS; (48)]. Backtracked RNA is shown as atomic spheres with yellow carbon atoms.

The RdRp NTP-entry tunnel provides a steric and electrostatic environment conducive to channeling the backtracked RNA out of the active site without specific polar protein-RNA interactions that could hinder the RNA movement (Fig. 3C and 4). Comparing the electrostatic surface potential of the NTP-entry tunnels of the SARS-CoV-2 RdRp with eukaryotic and bacterial DdRps reveals a similar overall electrostatic surface environment that may facilitate backtracked RNA entry (Fig. 3C; SI Appendix, Fig. S4B), including a ’track’ of conserved positively-charged Arg and Lys residues of motif F (SARS-CoV-2 nsp12 K545, K551, R553, and R555; Fig. 4; SI Appendix, Fig. S4A). Conserved residues of RdRp motifs C and E complete the active-site/NTP-entry tunnel environment surrounding the backtracked RNA (Fig. 4; SI Appendix, Fig. S4A).

**Fig. 4.**
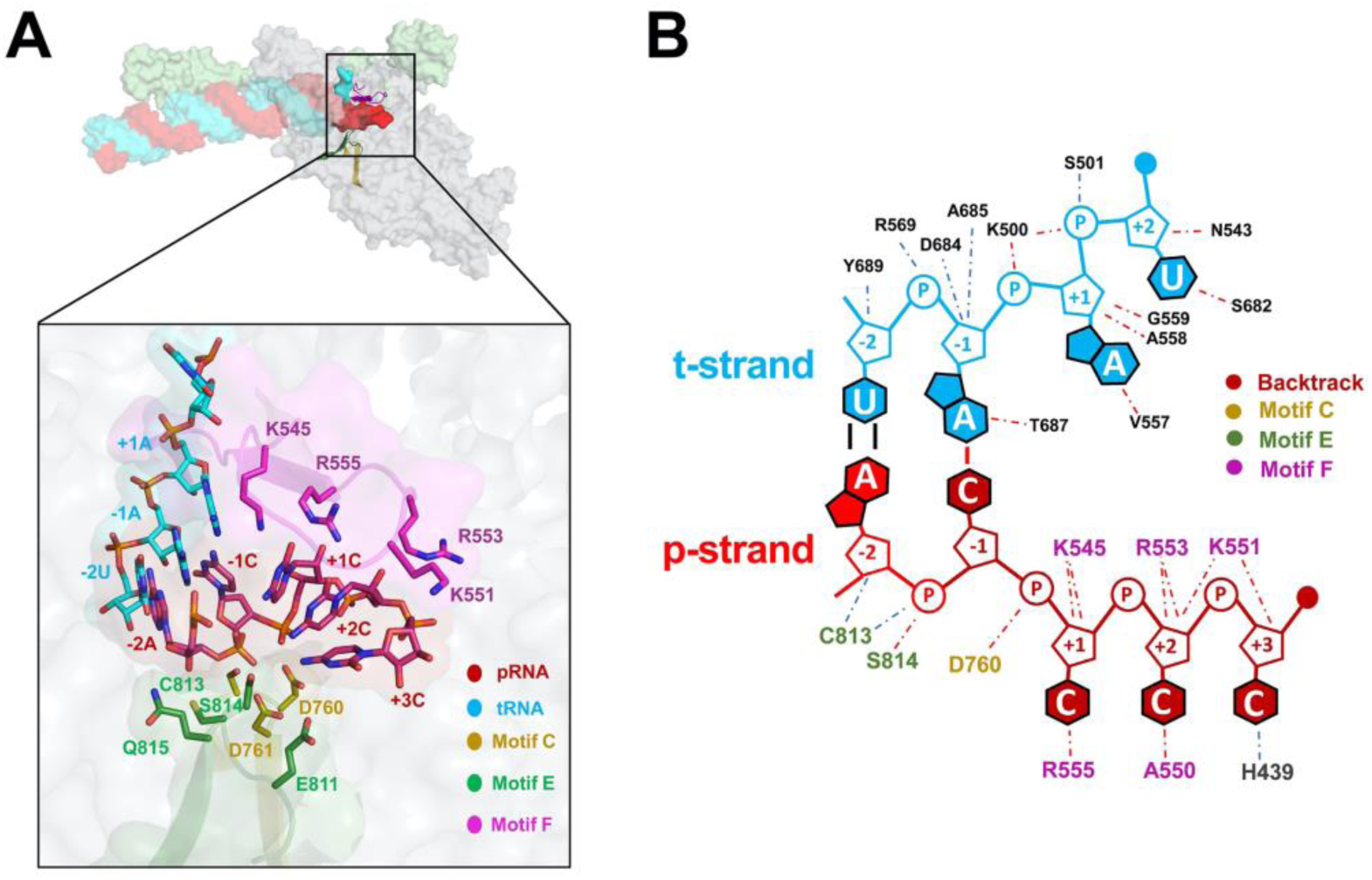
Protein-RNA interactions in the BTC. **A.** (*top*) Overall view of BTC_5_(local). Proteins are shown as transparent molecular surfaces, nucleic acids as atomic spheres. Nsp8a and nsp12 are shown (nsp7 and nsp8b are removed for clarity). Nsp12 motifs C, E, and F are shown as backbone ribbons (colored according to the key on the bottom. The boxed region is magnified below. (*bottom*) RNA is shown from -2 to +3. Proteins are shown as transparent molecular surfaces. RdRp motifs C, E, and F are shown as transparent backbone ribbons (colored according to the key) with side chains of residues that approach the backtracked RNA (≤ 4.5 Å) shown. **B.** Schematic illustrating the same protein-RNA interactions as (**A**). Drawn using Nucplot (49).

In the nsp13-RTCs, the RTC-scaffold (Fig. 1A) is bound in a post-translocated state (14); the 3’ p-RNA A is base-paired to the t-RNA U at the -1 site near the catalytic nsp12-D760 (Fig. 5A). The next t-RNA base (A at +1) is positioned to receive the incoming nucleoside-triphosphate (NTP) substrate, but the site for the incoming NTP substrate is empty (Fig. 5A). By contrast, the BTC structures were translocated by one base pair compared to the RTCs; the base pair corresponding to the A-U Watson-Crick base pair at the 3’-end of the p-RNA (located in the -1 site of the RTCs) was in the -2 position of the BTCs (Fig. 1A, 4, and 5B). The -1 position of the BTC was occupied by the first C-A mismatch; the p-RNA -1C made a non-Watson-Crick hydrogen bond with the opposing t-RNA A (Fig. 4 and 5B). The next three mismatched p-RNA nucleotides (+1C, +2C, +3C) trailed into the NTP entry tunnel (Fig. 4 and 5B). The 3’-nucleotide of the BTC_5_- scaffold p-RNA (+4C; Fig. 1A) was solvent-exposed at the outward-facing end of the NTP-entry tunnel, lacked density and was therefore not modelled (Fig. 2B). The trajectory of the backtracked nucleotides at positions +1/+2 was sharply bent due to spatial constraints of motif F residues (Fig. 4A).

**Fig. 5.**
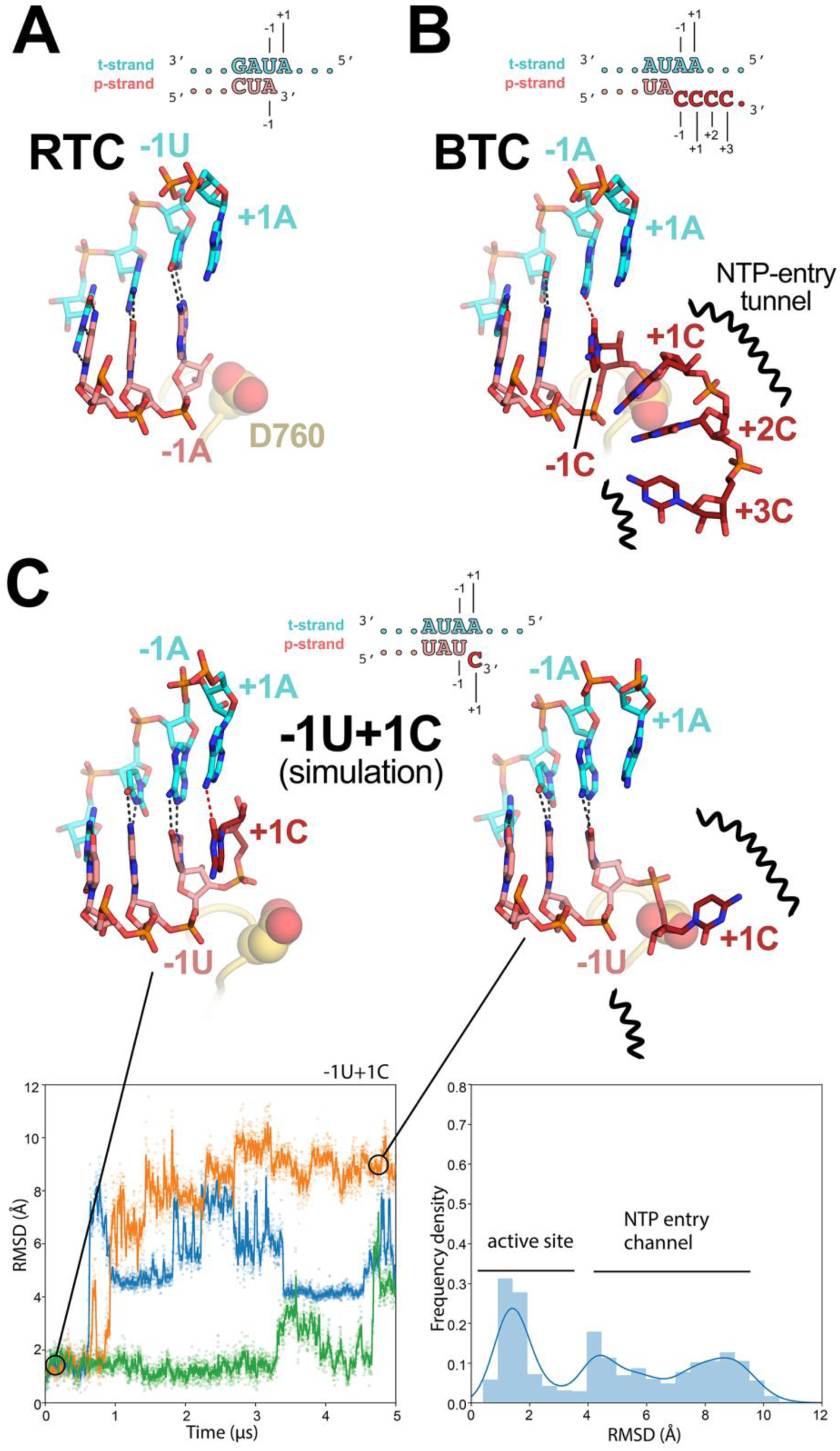
Comparison of active-site proximal RNA in the RTC and BTC structures, and from simulations of a mismatched nucleotide at the p-RNA 3’-end. **A-B,** Comparison of the active-site proximal RNA in the RTC [**A**; PDB ID: 6XEZ; (14)], BTC_5_(local) (**B**), and from selected snapshots of molecular dynamics simulations of a -1U+1C complex (**C**). The schematics denote the nucleotides shown in the context of the RTC- (**A**) and BTC_5_-scaffolds (**B**; full scaffold sequences shown in Figure 1A) or generated from the BTC_5_-scaffold for the simulations (**C**). Carbon atoms of the t-RNA are colored cyan, p-RNA are colored salmon except in the case of mismatched C’s at the 3’-end, which are colored dark red. Watson-Crick base pairing hydrogen-bonds are denoted as dark gray dashed lines, other hydrogen-bonds as red dashed lines. Nsp12 motif C is shown as a yellow-orange backbone ribbon, and the side-chain of D760 is shown as atomic spheres. **A.** The RTC is in a post-translocated state, with the A-U base pair at the p-RNA 3’-end in the -1 position (14). **B.** The BTC_5_(local) RNA is translocated compared to the RTC; the base pair corresponding to A-U at the 3’-end of the RTC RNA in the -1 position is in the -2 position of the BTC RNA. A C-A mismatch occupies the BTC -1 site. The +1, +2 and +3 mismatched C’s trail into the RdRp NTP-entry tunnel (denoted by black squiggly lines). The +4C (present in the BTC_5_-scaffold; Figure 1A) is exposed to solvent, disordered and not modelled. **C.** Molecular dynamics simulations of the nsp13_2_-BTC_−1U+1C_ complex. The complex was simulated with 3 replicates. RMSD values plotted as a function of time represent the heavy-atom RMSD of the +1C of the p-RNA compared with the starting configuration (see Methods). The RMSD histograms (plotted on the right) are an aggregate of all three replicates. Two structures taken from one of the simulations are shown, one showing the +1C of the p-RNA in the active site (*t* = 0 μs) and the other showing the +1C frayed into the NTP-entry tunnel (*t* = 4.5 μs).

### Nsp13 stimulates backtracking

The SARS-CoV-2 wild-type holo-RdRp required the nsp13 helicase to bind the BTC-scaffolds efficiently (Fig. 1B). However, we observed that the holo-RdRp containing nsp12 with a single amino acid substitution (D760A) did not require nsp13 to bind the BTC-scaffolds (SI Appendix; Fig. S1A, lane 4). Nsp12-D760 is a conserved residue of the RdRp motif C that chelates a crucial Mg^2+^ ion in catalytic complexes [SI Appendix; Fig. S4A; (32)], but in RdRp structures lacking substrate (including the BTC structures), the Mg^2+^ ions are absent (14, 36, 37). The catalytic Asp residues of the DdRps typically chelate the Mg^2+^ ion even in the absence of substrate (31, 38), and this Mg^2+^ is retained in DdRp backtracked structures (27–30). Our RdRp BTC structures suggest that in the absence of a Mg^2+^ ion, D760 presents an electrostatic barrier to the phosphate backbone of the backtracked RNA (Fig. 5B), explaining the requirement for the helicase to surmount this barrier and why removal of D760 stabilizes binding to the BTC-scaffolds.

To generate the SARS-CoV-2 BTCs for structural studies, we used the BTC_5_-scaffold with five mismatched C’s at the p-RNA 3’-end (Fig. 1A). To study the formation of SARS-CoV-2 BTCs from an RTC-scaffold (fully Watson-Crick base paired p-RNA 3’-end), we analyzed UV-induced crosslinking from 4-thio-U incorporated penultimate to the p-RNA 3’-end [RTC(4-thio-U)-scaffold; Si Appendix, Fig. S5A; (39)]. Crosslinking was absolutely dependent on the presence of 4-thio-U in the RNA, establishing specificity (SI Appendix; Fig. S5B). RTCs assembled with wild-type nsp12 and the RTC(4-thio-U)-scaffold gave little to no protein-RNA crosslinking upon UV exposure (SI Appendix; Fig. S5A, lane 2). These conditions favor a post-translocated RTC (14, 36, 37) with the 4-thio-U sequestered in the RNA-RNA hybrid and thus not available for protein-RNA crosslinking. Crosslinking of the p-RNA to nsp12 was substantially increased by the addition of nsp13 (with 2 mM ATP, which is present in all the lanes; SI Appendix; Fig. S5A, lane 3). Under these conditions, we propose that the translocation activity of nsp13 backtracked a fraction of the complexes, freeing the 4-thio-U from the RNA-RNA hybrid for crosslinking to nsp12. Replacing wild-type nsp12 with nsp12-D760A (nsp12*; SI Appendix, Fig. S5A, lanes 6-8), which is more prone to backtracking (SI Appendix; Fig. S1A), increased the protein-RNA crosslinking under all conditions, with the maximal crosslinking occurring under the conditions expected to favor backtracking the most (SI Appendix; Fig. S5A, lane 7). These results affirm the view that nsp13 facilitates backtracking of the SARS-CoV-2 RdRp.

### A mismatched nucleotide at the p-RNA 3’-end spontaneously frays and enters into the RdRp NTP-entry tunnel

The SARS-CoV-2 RTC is a highly processive and rapid replicase/transciptase, capable of replicating an ∼1 kb RNA template at an average rate of ∼170 nt/s (40). However, studies of other viral RdRps suggest that misincorporation slows the overall elongation rate and may induce backtracking (41–43). We used molecular dynamics simulations to explore the fate of a mismatched nucleotide incorporated at the p-RNA 3’-end. Starting with the nsp13_2_-BTC_5_ structure, the -1C was mutated to U, and the +2 to +4 C’s were removed. The resulting pre-translocated p-RNA had a matched -1U and a mismatched +1C (-1U+1C; Fig. 5C). In three 5 μs simulations we observed the 3’-mismatched +1C alternating between two positions, either remaining in the vicinity of the active site (RMSD < 3.5 Å) or fraying away from the p-RNA:t-RNA hybrid towards or into the NTP-entry tunnel (RMSD > 3.5 Å; Fig. 5C). Based on analysis of the aggregated -1U+1C simulations, the mismatched +1C spent about 40% of the time near the active site and about 60% of the time frayed towards or in the NTP-entry tunnel. In control simulations with a fully matched p-RNA 3’-end (-1U+1U), the matched +1U at the p-RNA 3’-end did not fray and spent 100% of the time in the active site pocket (SI Appendix; Fig. S6).

## Discussion

Our results establish that the SARS-CoV-2 RTC backtracks, that backtracking is facilitated by the nsp13 helicase, and that the resulting single-stranded 3’- segment of the p-RNA extrudes out the RdRp NTP-entry tunnel in a manner analogous to the evolutionarily unrelated cellular DdRps (Fig. 3). Thus, a secondary tunnel to accommodate backtracked RNA, facilitating fidelity and possibly other functions (Fig. 6), appears to be a crucial feature of transcriptase enzymes that evolved independently.

**Fig. 6.**
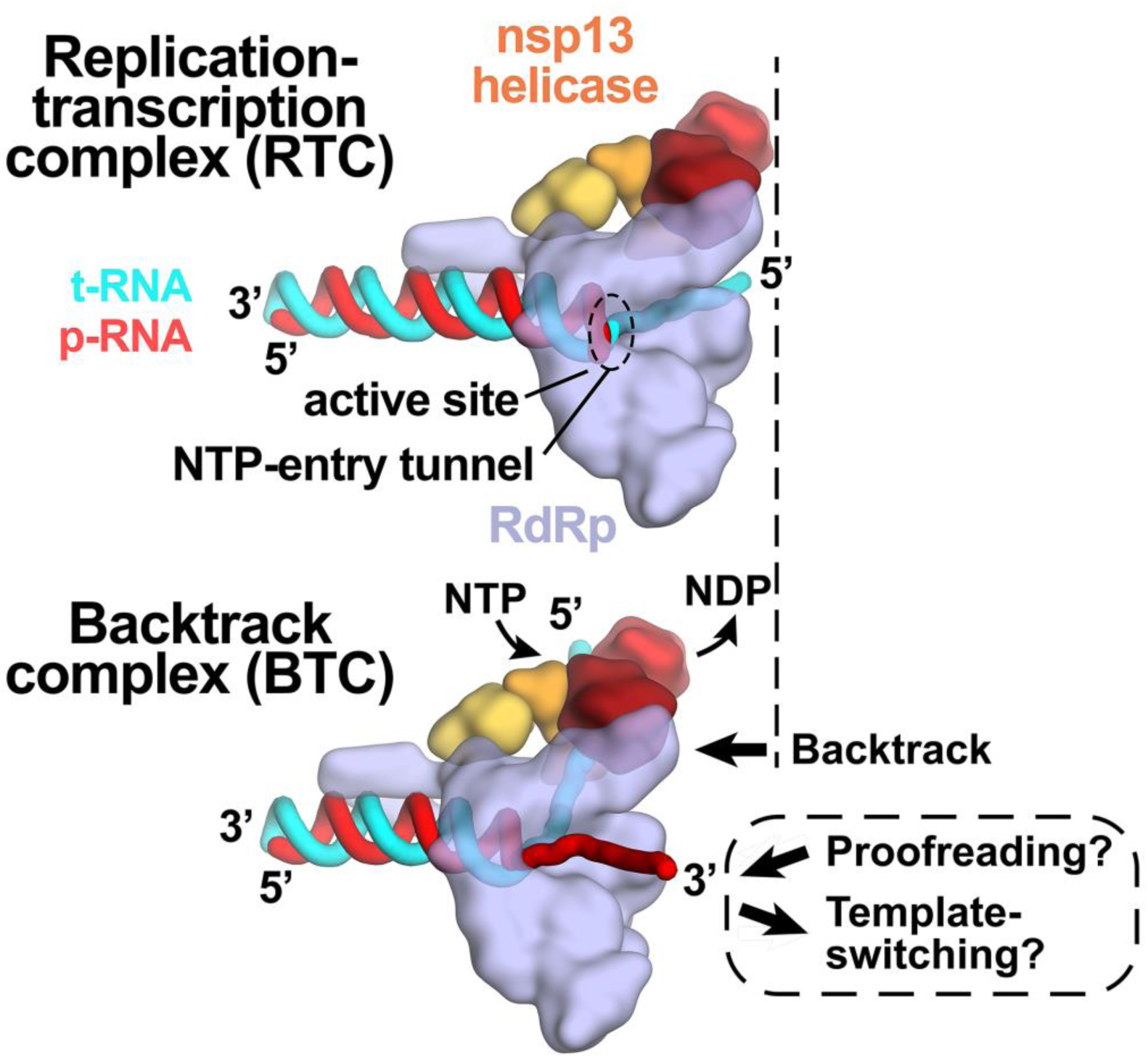
Role of backtracking in proofreading and template-switching during sub-genomic transcription. Schematic illustrating the proposed model for backtracking of the SARS-CoV-2 RTC and its potential role in proofreading and template-switching during sub-genomic transcription. The structural models are shown as cartoons (holo-RdRp, light blue; nsp13 helicase, orange shades; RNA strands, colored tubes as indicated). (*top*) In the RTC, the elongating RdRp moves from left-to-right. The RdRp active site holds the p-RNA 3’-end. The NTP-entry tunnel provides access from solution to the RdRp active site. The downstream (5’) single-stranded t-RNA is not engaged with nsp13. (*bottom*) In the BTC, nsp13 translocates on the downstream (5’) single-stranded t-RNA, pushing the RdRp backwards (right-to-left) on the RNA. This causes the p-RNA to reverse-thread through the complex, with the resulting single-stranded 3’-fragment extruding out the NTP-entry tunnel. The exposure of the p-RNA 3’- end could facilitate proofreading (9, 10, 12, 50) and also template-switching during sub-genomic transcription (7, 34).

Backtracking of Φ6 and poliovirus RdRps has been reported based on analysis of single-molecule observations (41–43). The nsp13 helicase facilitates efficient backtracking of the SARS-CoV-2 RTC (SI Appendix; Fig. S5). We note that in bacteria, the UvrD helicase has been shown to induce DdRp backtracking, suggesting that a role for helicases in backtracking may be widespread (44).

Our results are consistent with the view that a matched nucleotide at the pre-translocated p-RNA 3’-end remains base paired to the t-RNA (Fig. 5; SI Appendix, Fig. S6), facilitating translocation and subsequent NTP addition and thus rapid elongation (at a maximum elongation rate of ∼170 nt/s, a translocation event would occur approximately every 6 msec, on average, explaining why translocation was not observed in our 5 μs simulations; Fig. 5; SI Appendix, Fig. S6). However, upon misincorporation, the pre-translocated, mismatched nucleotide at the p-RNA 3’-end spends more than half the time frayed from the t-RNA and towards or in the NTP-entry tunnel (Fig. 5C), a configuration that is likely recalcitrant to translocation and subsequent elongation. The favorable environment of the NTP-entry tunnel (Fig. 3 and 4) may further encourage backtracking. The resulting inhibition of translocation may enable the tight engagement of the nsp13.1 helicase with the downstream single-stranded t-RNA (Fig. 2A), allowing the 5’->3’ translocation activity of the helicase to more robustly backtrack the complex (SI Appendix; Fig. S5).

Our findings have implications for the processes of subgenomic transcription and proofreading in SARS-CoV-2 [Fig. 6; (14)]. Generation of mRNAs for the viral structural proteins begins with transcription initiation at the 3’-poly(A) tail of the (+)-strand RNA genome. The process, called sub-genomic transcription, ultimately generates a nested set of transcripts that are both 5’- and 3’-co-terminal with the viral genome and involves a remarkable template-switch from the 3’-portion of the genome to the 5’-leader (7, 34). The template-switching event is thought to involve base-pairing between the 3’-end of the nascent transcript and a complementary sequence (the Transcription Regulatory Sequence, or TRS) near the (+)-strand 5’-leader (45). Backtracking could extrude the 3’-end of the nascent transcript out the NTP-entry tunnel, making it available for base pairing to the 5’-TRS (Fig. 6). Our results establishing that the SARS-CoV-2 RTC can backtrack validates a key prediction of this model for the mechanism of template-switching during sub-genomic transcription (14).

Nucleotide analogs that function by being incorporated into product RNA by viral RdRps are important antiviral therapeutics (46). Notably, their incorporation may induce backtracking by the RdRp (43). Rdv, a nucleotide analog, is the only FDA-approved drug for COVID-19 treatment (5). Our results support a model in which RdRp misincorporation or incorporation of nucleotide analogs can pause the RdRp, allowing nsp13 to engage with the downstream single-stranded t-RNA to induce backtracking (14). The resulting exposure of the p-RNA 3’-end out the NTP-entry tunnel (Fig. 3A and 6) could provide access for the SARS-CoV-2 proofreading machinery [nsp10/14; (9, 12)] to degrade the p-RNA 3’-end, thus removing the misincorporation or analog. This proofreading activity, which is unique to the nidovirus order to which CoVs belong (10), is a major determinant for the resistance of CoVs against many nucleotide analog inhibitors (13). Thus, understanding RdRp backtracking and its potential role in CoV proofreading can facilitate the development of therapeutics.

## Materials and Methods

Detailed descriptions of SARS-CoV-2 nsp12, 7, 8, and 13 protein purification, assembly of the RTC complexes, Native EMSAs, native mass-spectrometry, cross-linking, specimen preparation for cryo-EM, cryo-EM data acquisition and processing, model building and refinement, and molecular dynamics simulations are provided in the SI Appendix.

## Acknowledgments

We thank M. Ebrahim, J. Sotiris, and H. Ng at The Rockefeller University Evelyn Gruss Lipper Cryo-electron Microscopy Resource Center for help with cryo-EM. Some of the work reported here was conducted at the Simons Electron Microscopy Center (SEMC) and the National Resource for Automated Molecular Microscopy (NRAMM) and National Center for CryoEM Access and Training (NCCAT) located at the NYSBC, supported by grants from the NIH National Institute of General Medical Sciences (P41 GM103310), NYSTAR, the Simons Foundation (SF349247), the NIH Common Fund Transformative High Resolution Cryo-Electron Microscopy program (U24 GM129539) and NY State Assembly Majority. This work was supported by the Pels Family Center for Biochemistry and Structural Biology (The Rockefeller University), and NIH grants P41 GM109824 and P41 GM103314 to B.T.C, R35 GM118130 to S.A.D, and R01 GM114450 to E.A.C.

## Supplementary Information Text

### METHODS

Structural biology software was accessed through the SBGrid consortium (1).

#### Protein expression and purification

*SARS-CoV-2 nsp12*. SARS-CoV-2 nsp12 was expressed and purified as described (1). A pRSFDuet-1 plasmid expressing SARS-CoV-2 His_6_-SUMO-nsp12 (Addgene plasmid 159107) was transformed into *Escherichia coli* (*Eco*) BL21-CodonPlus cells (Agilent). Cells were grown, followed by the addition of isopropyl β-d-1-thiogalactopyranoside (IPTG) to induce protein expression overnight. Cells were collected by centrifugation, resuspended and lysed in a continuous-flow French press (Avestin). The lysate was cleared by centrifugation, loaded onto a HiTrap Heparin HP column (Cytiva), and then eluted using a salt gradient. The fractions containing nsp12 were pooled and loaded onto a HisTrap HP column (Cytiva), washed, and eluted. Eluted nsp12 was dialyzed overnight in the presence of His_6_-Ulp1 SUMO protease. Cleaved nsp12 was passed through a HisTrap HP column (Cytiva). Flow-through was collected, concentrated by centrifugal filtration (Amicon), and loaded on a Superdex 200 Hiload 16/600 (Cytiva) for size-exclusion chromatography. Glycerol was added to the purified nsp12, aliquoted, flash frozen with liquid N_2_, and stored at -80°C.

*SARS-CoV-2 nsp7/8*. SARS-CoV-2 nsp7/8 was expressed and purified as described (1). The pCDFDuet-1 plasmid expressing SARS-CoV-2 His_6_-ppx-nsp7/8 (ppx is a Prescission Protease cleavage site; Addgene plasmid 159092) was transformed into *Eco* BL21(DE3). Cells were grown and protein expression was induced overnight by the addition of IPTG. Cells were collected by centrifugation, resuspended, and lysed in a continuous-flow French press (Avestin). The lysate was cleared by centrifugation, then loaded onto a HisTrap HP column (Cytiva), washed, and eluted. Eluted nsp7/8 was dialyzed overnight in the presence of His_6_-Prescission Protease to cleave the His_6_-tag. Cleaved nsp7/8 was passed through a HisTrap HP column (Cytiva). Flow-through was collected, concentrated by centrifugal filtration (Amicon), and loaded onto a Superdex 75 Hiload 16/600 (Cytiva). Glycerol was added to the purified nsp7/8, aliquoted, flash frozen with liquid N_2_, and stored at -80°C.

*SARS-CoV-2 nsp13*. SARS-CoV-2 nsp13 was expressed and purified as described (1). The pet28 plasmid containing SARS-CoV-2 His_6_-ppx-nsp13 (Addgene plasmid 159390) was transformed into *Eco* Rosetta(DE3) (Novagen). Cells were grown, followed by the addition of IPTG to induce protein expression overnight. Cells were collected by centrifugation, resuspended, and lysed in a continuous-flow French press (Avestin). The lysate was cleared by centrifugation, then loaded onto a HisTrap HP column (Cytiva), washed, and eluted. Eluted nsp13 was dialyzed overnight in the presence of His_6_-Prescission Protease to cleave His_6_-tag. Cleaved nsp13 was passed through a HisTrap HP column (Cytiva). Flow-through was collected, concentrated by centrifugal filtration (Amicon), and loaded onto a Superdex 200 Hiload 16/600 (Cytiva). Glycerol was added to the purified nsp13, aliquoted, flash frozen with liquid N_2_, and stored at - 80°C.

#### Native electrophoretic mobility shift assays

Nsp12 or nsp12-D760A were incubated with 3-fold molar excess of nsp7/8 in transcription buffer (120 mM K-acetate, 20 mM HEPES pH 8, 10 mM MgCl_2_, 2 mM DTT) to assemble holo-RdRp (2 μM final). The resulting complex was incubated with 1 μM of annealed RNA scaffold (Horizon Discovery) for 5 minutes at 30°C. Nsp13 and pre-mixed ADP and AlF_3_ (Sigma-Aldrich) were added to a final concentration of 2 μM and 2 mM, respectively, and incubated for an additional 5 minutes at 30°C. Reactions were analyzed by native gel electrophoresis on a 4.5% polyacrylamide native gel (37.5:1 acrylamide:bis-acrylamide) in 1X TBE (89 mM Tris, 89 mM boric acid, 1 mM EDTA) at 4°C. The gel was stained with Gel-Red (Biotium).

#### Native mass spectrometry (nMS) analysis

The reconstituted sample containing 4 µM RTC and 8 µM nsp13 incubated with 2 mM ADP-AlF_3_ was buffer-exchanged into 150 mM ammonium acetate, 0.01% Tween-20, pH 7.5 using a Zeba microspin desalting column with a 40 kDa MWCO (ThermoFisher Scientific). For nMS analysis, a 2–3 µL aliquot of the buffer-exchanged sample was loaded into a gold-coated quartz capillary tip that was prepared in-house and then electrosprayed into an Exactive Plus with extended mass range (EMR) instrument (Thermo Fisher Scientific) with a static direct infusion nanospray source (2). The MS parameters used: spray voltage, 1.2 kV; capillary temperature, 150 °C; in-source dissociation, 0 V; S-lens RF level, 200; resolving power, 17,500 at *m/z* of 200; AGC target, 1 x 10^6^; maximum injection time, 200 ms; number of microscans, 5; injection flatapole, 6 V; interflatapole, 4 V; bent flatapole, 4 V; high energy collision dissociation (HCD), 200 V; ultrahigh vacuum pressure, 7.2 × 10^−10^ mbar; total number of scans, at least 100. Mass calibration in positive EMR mode was performed using cesium iodide. For data processing, the acquired MS spectra were visualized using Thermo Xcalibur Qual Browser (v. 4.2.47). MS spectra deconvolution was performed either manually or using the software UniDec v. 4.2.0 (3, 4). The following parameters were used for the UniDec processing: m/z range, 7,000 – 10,000 Th; background subtraction, subtract curved at 100; smooth charge state distribution, enabled; peak shape function, Gaussian; Beta Softmax distribution parameter, 20.

The expected masses for the component proteins based on previous nMS experiments (1) include nsp7: 9,137 Da; nsp8 (N-terminal Met lost): 21,881 Da; nsp13 (post-protease cleavage, has three Zn^2+^ ions coordinated with 9 deprotonated cysteine residues): 67,464 Da, and nsp12 (has two Zn^2+^ ions coordinated with 6 deprotonated cysteine residues): 106,785 Da. The mass of the assembled RNA duplex scaffold is 30,512 Da.

Experimental masses were reported as the average mass ± standard deviation (S.D.) across all the calculated mass values within the observed charge state series. Mass accuracies were calculated as the % difference between the measured and expected masses relative to the expected mass. The observed mass accuracies ranged from 0.016 – 0.035%.

#### Preparation of SARS-CoV-2 nsp13-BTC_5_ for Cryo-EM

Purified nsp12 and nsp7/8 were mixed in a 1:3 molar ratio and incubated at 22° C for 15 minutes. The mixture was buffer-exchanged into cryo-EM buffer (20 mM HEPES pH 8.0, 150 mM K-acetate, 10 mM MgCl_2_, 1 mM DTT) using Zeba desalting columns (ThermoFisher Scientific) and incubated with annealed BTC_5_-scaffold (Fig. 1A) in a 1:1.5 molar ratio. Purified nsp13 was concentrated by centrifugal filtration (Amicon) and buffer exchanged into cryo-EM buffer using Zeba desalting columns. Buffer exchanged nsp13 was mixed with ADP and AlF_3_ and then added to nsp7/8/12/RNA scaffold at a molar ratio of 1:1 with a final concentration of 2 mM ADP-AlF_3_. Complex was incubated for 5 minutes at 30° C and further concentrated by centrifugal filtration (Amicon).

#### Cryo-EM grid preparation

Prior to grid freezing, 3-([3- cholamidopropyl]dimethylammonio)-2-hydroxy-1-propanesulfonate (CHAPSO, Anatrace) was added to the sample (8 mM final), resulting in a final complex concentration of 10 µM. The final buffer condition for the cryo-EM sample was 20 mM HEPES pH 8.0, 150 mM K-acetate, 10 mM MgCl_2_, 1 mM DTT, 2 mM ADP-AlF_3_, 8 mM CHAPSO. C-flat holey carbon grids (CF-1.2/1.3-4Au, Electron Microscopy Sciences) were glow-discharged for 20 s prior to the application of 3.5 μL of sample. Using a Vitrobot Mark IV (ThermoFisher Scientific), grids were blotted and plunge-frozen into liquid ethane with 90% chamber humidity at 4°C.

#### Cryo-EM data acquisition and processing

Structural biology software was accessed through the SBGrid consortium (5). Grids were imaged using a 300 kV Titan Krios (ThermoFisher Scientific) equipped with a K3 camera (Gatan) and a BioQuantum imaging filter (Gatan). Images were recorded using Leginon (6) with a pixel size of 1.065 Å/px (micrograph dimensions of 5,760 x 4,092 px) over a nominal defocus range of -0.8 μm to -2.5 μm and 30 eV slit width. Movies were recorded in “counting mode” (native K3 camera binning 2) with ∼30 e-/px/s in dose-fractionation mode with subframes of 50 ms over a 2.5 s exposure (50 frames) to give a total dose of ∼66 e-/Å. Dose-fractionated movies were gain-normalized, drift-corrected, summed, and dose-weighted using MotionCor2 (7). The contrast transfer function (CTF) was estimated for each summed image using the Patch CTF module in cryoSPARC v2.15.0 (8) Particles were picked and extracted from the dose-weighted images with box size of 256 px using cryoSPARC Blob Picker and Particle Extraction. The entire dataset consisted of 10,685 motion-corrected images with 4,961,691 particles. Particles were sorted using cryoSPARC 2D classification (N=100), resulting in 2,412,034 curated particles. Initial models (Ref 1: decoy 1, Ref 2: complex, Ref 3: decoy 2; SI Appendix; Fig. S2) were generated using cryoSPARC *ab initio* Reconstruction on a subset of 85,398 particles. Particles were further curated using Ref 1-3 as 3D templates for cryoSPARC Heterogeneous Refinement (N=6), resulting in the following: class1 (Ref 1), 258,097 particles; class2 (Ref 1), 263,966 particles; class3 (Ref 2), 668,743 particles; class4 (Ref 2), 665,480 particles; class5 (Ref 3), 280,933 particles; class6 (Ref 3), 274,815 particles. Particles from class3 and class4 were combined and further curated with another round of Heterogeneous Refinement (N=6), resulting in the following: class1 (Ref 1), 67,639 particles; class2 (Ref 1), 61,097 particles; class3 (Ref 2), 553,368 particles; class4 (Ref 2), 554,581 particles; class5 (Ref 3), 42,114 particles; class6 (Ref 3), 55,424 particles. Curated particles from class3 and class4 were combined, re-extracted with a box size of 320 px, and further classified using Ref 2 as a 3D template for cryoSPARC Heterogeneous Refinement (N=4). Classes from this round of Heterogeneous Refinement (N=4) were as follows: class1 (Ref 2), 871,163 particles; class2 (Ref 2), 77,769 particles; class3 (Ref 2), 61,489 particles; class4 (Ref 2), 64,026 particles. Particles from class1 and class2 were combined and further sorted using Heterogeneous Refinement (N=4) using class maps as templates, resulting in the following: class1, 134,536 particles; class2, 270,170 particles; class3, 294,162 particles; class4, 172,295 particles. Classification revealed two unique classes: nsp13_1_-BTC (class1 and class2) and nsp13_2_-BTC (class3 and class4). Particles within each class were further processed using RELION 3.1-beta Bayesian Polishing(9, 10). Polished particles were refined using cryoSPARC Non-uniform Refinement, resulting in structures with the following particle counts and nominal resolutions: nsp13_1_-BTC (404,706 particles; 3.40 Å) and nsp13_2_-BTC (466,457 particles; 3.45 Å).

To improve the resolution of the RNA in the BTC, particles from both classes were combined in a cryoSPARC Non-uniform Refinement and density corresponding to nsp13 was subtracted. Subtracted particles were further refined with cryoSPARC Local Refinement using a mask encompassing the BTC and a fulcrum point defined on the backtracked RNA. This map, BTC_5_(local), contained 871,163 particles with a nominal resolution of 3.23 Å.

To improve the density of nsp13.2 in the nsp13_2_-BTC map, particles were subtracted using a mask defined around nsp13.2, leaving residual signal for only nsp13.2. Subtracted particles were classified (N=4) in RELION 3.1 beta using a mask around nsp13.2, resulting in the following classes: class1, 71,607 particles; class2, 163,540 particles; class3, 176,461 particles; class4, 54,849 particles. Subtracted particles in class1 and class2 were combined and reverted back to the original particles, followed by refinement using cryoSPARC Non-uniform Refinement. The resulting map of nsp13_2_-BTC contains 235,147 particles with nominal resolution of 3.59 Å. Local resolution calculations were generated using blocres and blocfilt from the Bsoft package (11).

#### Model building and refinement

Initial models were derived from PDB: 6XEZ (1). The models were manually fit into the cryo-EM density maps using Chimera (12) and rigid-body and real-space refined using Phenix real_space_refine (13). For real-space refinement, rigid body refinement was followed by all-atom and B-factor refinement with Ramachandran and secondary structure restraints. Models were inspected and modified in Coot (14).

#### 4-thiouridine crosslinking

Nsp12 or nsp12-D760A were incubated with 3-fold molar excess of nsp7/8 to assemble holo-RdRp (2 μM final) in transcription buffer. The resulting holo-RdRp was added to a modified RNA scaffold (SI Appendix; Fig. S5A) containing a photoactivable 4-thiouridine base (Horizon Discovery) which was 5’-labelled by T4-polynucleotide kinase (New England Biolabs) with γ-^32^P-ATP (Perkin-Elmer). The holo-RdRp/RNA complex was left to incubate for 5 minutes at 30°C in the dark. Nsp13 and ATP were added to a final concentration of 2 uM and 2 mM, respectively, and incubated for five minutes at 30°C in the dark. The reaction mixture was transferred to a Parafilm covered aluminum block at 4°C and irradiated with a 365-nm handheld UV lamp. Reactions were quenched with LDS sample loading buffer (ThermoFisher Scientific) and analyzed by gel electrophoresis on a NuPAGE 4-12% Bis-Tris gel (ThermoFisher) at 150 Volts for 1 hour and visualized by autoradioagraphy.

#### Molecular dynamics simulations

##### General simulation setup and parameterization

Proteins, ADP, and ions were parameterized with the DES-Amber SF1.0 force field (15). RNAs were parameterized with the Amber ff14 RNA force field (16) with modified electrostatic, van der Waals, and torsional parameters to more accurately reproduce the energetics of nucleobase stacking (17). The systems were solvated with water parameterized with the TIP4P-D water model (18) and neutralized with 150 mM NaCl buffer. The systems each contained ∼887,000 atoms in a 190×190×190 Å cubic box.

Systems were first equilibrated on GPU Desmond using a mixed NVT/NPT schedule (19), followed by a 1 µs relaxation simulation on Anton, a special-purpose machine for molecular dynamics simulations (20). All production simulations were performed on Anton and initiated from the last frame of the relaxation simulation. Production simulations were performed in the NPT ensemble at 310 K using the Martyna-Tobias-Klein barostat (21). The simulation time step was 2.5 fs, and a modified r-RESPA integrator (22, 23) was used in which long-range electrostatic interactions were evaluated every three time steps. Electrostatic forces were calculated using the *u*-series method (24). A 9-Å cutoff was applied for the van der Waals calculations.

##### System preparation

The nsp13_2_-BTC_-1U+1C_ and the nsp13_2_-BTC_-1U+1U_ complexes were prepared from the cryo-EM structure of the nsp13_2_-BTC_5._ AlF_3_ and CHAPSO were removed. Cytosines at the +2 and +3 positions of the p-RNA were removed, and the cytosine at −1 was mutated to uracil. The resulting p-RNA had a matched −1U and a mismatched +1C in nsp13_2_-BTC_−1U+1C_, and a matched −1U and +1U in nsp13_2_-BTC_−1U+1U_. Missing loops and termini in proteins were capped with ACE/NME capping groups. The two complexes were prepared for simulation using the Protein Preparation Wizard in Schrödinger Maestro. After a 1 µs relaxation simulation of the nsp13_2_-BTC_−1U+1C_ complex, the −1U of the p-RNA formed a Watson-Crick base pair with the −1A in the t-RNA, and the +1C of p-RNA formed a non-Watson-Crick C-A hydrogen bond with the +1A of the t-RNA in the active site. After a 1 µs relaxation simulation of the nsp13_2_-BTC_−1U+1U_ complex, the −1U and +1U of the p-RNA formed Watson-Crick base pairs with the −1A and +1A of the t-RNA respectively.

##### Simulation analysis

All simulations were visually inspected using the in-house visualization software Firefly. The average root-mean-square deviation (RMSD) was calculated for +1C (or +1U) of the p-RNA between the last frame of the 1 µs relaxation simulation and instantaneous structures from the trajectories, aligned on the entire nps12 module.

#### Quantification and statistical analysis

The nMS spectra were visualized using Thermo Xcalibur Qual Browser (versions 3.0.63 and 4.2.27), deconvolved using UniDec versions 3.2 and 4.1 (3, 4) and plotted using the m/z software (Proteometrics LLC, New York, NY). Experimental masses (SI Appendix; Fig. S1B and C) were reported as the average mass ± standard deviation across all the calculated mass values obtained within the observed charge state distribution.

The local resolution of the cryo-EM maps (SI Appendix; Fig. S3B-D) was estimated using blocres (11) with the following parameters: box size 15, sampling 1.1, and cutoff 0.5. Directional 3D FSC (SI Appendix; Fig. S3H-J) were calculated by 3DFSC (25). The quantification and statistical analyses for model refinement and validation were generated using MolProbity (26) and PHENIX (13).

---

**Table S1.** Cryo-EM data collection, refinement, and validation statistics.

|  | **nsp7/8/12/13/BTC_scaffold/ADP-AIF<sub>3</sub>/CHAPSO** |  |  |
| --- | --- | --- | --- |
| **Sample ID** | nsp13<sub>1</sub>-BTC | nsp13<sub>2</sub>-BTC | BTC (local) |
| **EMDB** | EMD-23007 | EMD-23008 | EMD-23009 |
| **PDB** | 7KRN | 7KRO | 7KRP |
| **Data collection and processing** |  |  |  |
| Microscope |  | TFS Titan Krios |  |
| Voltage (kV) |  | 300 |  |
| Detector |  | Gatan K3 Camera |  |
| Electron exposure (e<sup>-</sup>/Å<sup>2</sup>) |  | 66 |  |
| Defocus range (μm) |  | -0.8 to -2.5 |  |
| Data collection mode |  | Counting Mode |  |
| Nominal Magnification |  | 81,000x |  |
| Pixel size (Å) |  | 1.065 |  |
| Symmetry imposed |  | C1 |  |
| Initial particle images (no.) |  | 4,961,691 |  |
| Final particle images (no.) | 404,706 | 235,147 | 871,163 |
| Map resolution (Å) - FSC threshold 0.143 | 3.40 | 3.59 | 3.23 |
| Map resolution range (Å) | 2.5-5.0 | 2.5-5.0 | 2.5-5.1 |
| **Refinement** |  |  |  |
| Initial model used (PDB code) | 6XEZ | 6XEZ | 6XEZ |
| Map sharpening B factor (Å<sup>2</sup>) | -139.6 | -127.6 | -103.9 |
| Model composition |  |  |  |
| Non-hydrogen atoms | 17351 | 21988 | 12561 |
| Protein residues | 1963 | 2553 | 1373 |
| Nucleic acid residues | 80 | 80 | 73 |
| Ligands | 5 Zn<sup>2+</sup>, 2 Mg<sup>2+</sup>, 3 CHAPSO,<br>2 ADP, 1 AlF<sub>3</sub> | 8 Zn<sup>2+</sup>, 3 Mg<sup>2+</sup>, 3 CHAPSO,<br>3 ADP, 2 AlF<sub>3</sub> | 2 Zn<sup>2+</sup>, 1 Mg<sup>2+</sup>, 3 CHAPSO,<br>1 ADP |
| B factors (Å<sup>2</sup>) |  |  |  |
| Protein | 45.65 | 74.56 | 38.31 |
| Nucleic acid | 128.79 | 163.6 | 140.77 |
| Ligands | 59.03 | 78.99 | 46.55 |
| R.m.s. deviations |  |  |  |
| Bond lengths (Å) | 0.007 | 0.007 | 0.004 |
| Bond angles (°) | 0.711 | 0.735 | 0.609 |
| Validation |  |  |  |
| MolProbity score | 2.66 | 2.68 | 1.97 |
| Clashscore | 9.18 | 8.96 | 6.12 |
| Poor rotamers (%) | 7.54 | 9.4 | 4.18 |
| Ramachandran plot |  |  |  |
| Favored (%) | 91.4 | 92.76 | 97.07 |
| Allowed (%) | 8.6 | 6.93 | 2.93 |
| Disallowed (%) | 0 | 0.31 | 0 |

## SUPPLEMENTAL FIGURES

**Fig. S1.**
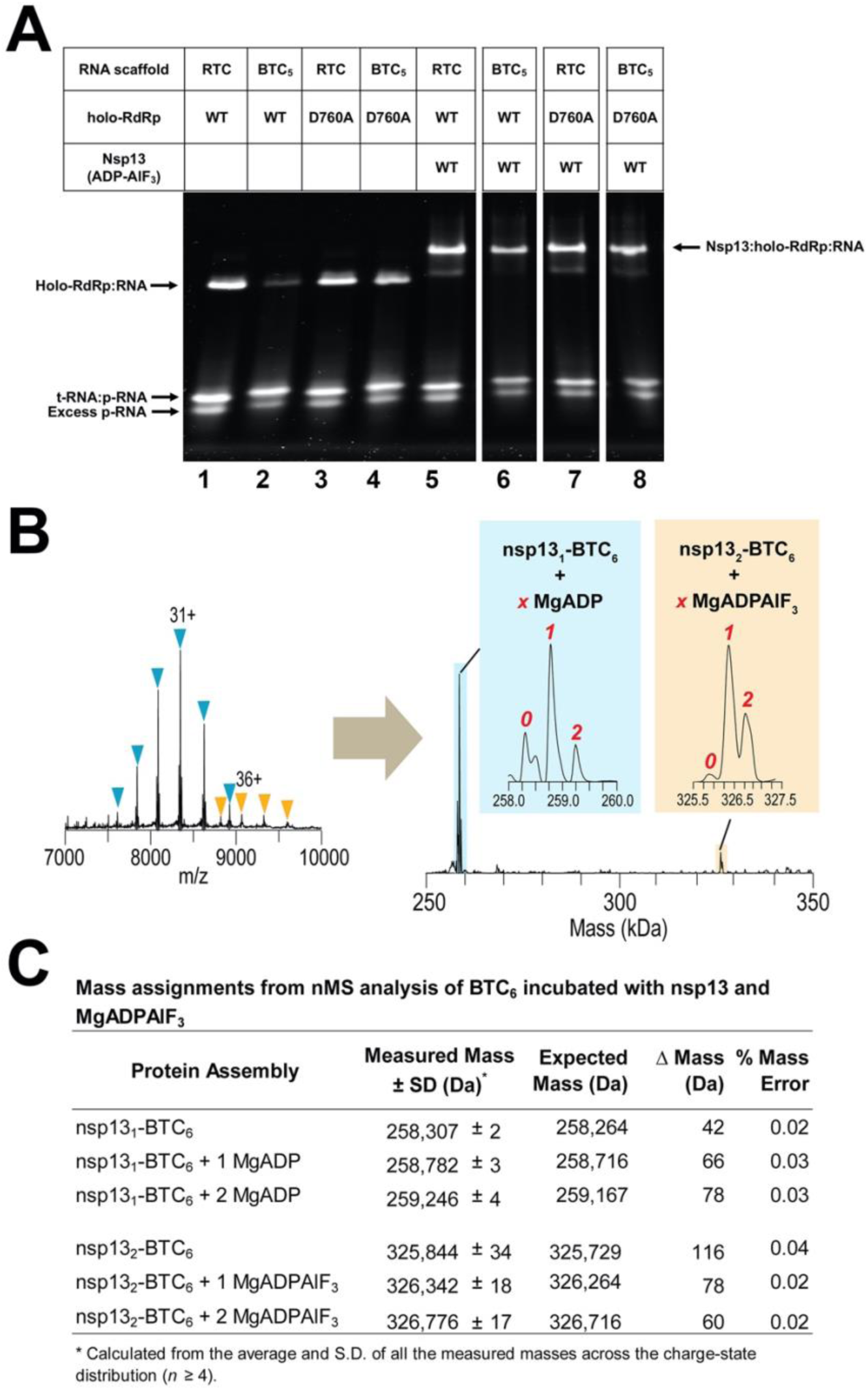
Native gel electrophoresis mobility shift assay and nMS analysis of the BTC. **A.** A native gel electrophoretic mobility shift assay reveals that wt-holo-RdRp requires nsp13(ADP-AlF_3_) to bind the BTC_5_-scaffold efficiently (compare lanes 1, 2, and 6) but holo-RdRp with nsp12-D760A does not require nsp13 (lane 4). **B.** The nMS spectrum and the deconvolved mass spectrum showing assembly of stable nsp13-BTC_6_ complexes. The peak for the nsp13_2_-BTC_6_ assembly is present at about ∼9% intensity relative to the predominant peak from nsp13_1_-BTC_6_. **C.** Mass assignments of the deconvolved peaks from the nMS analysis.

**Fig. S2.**
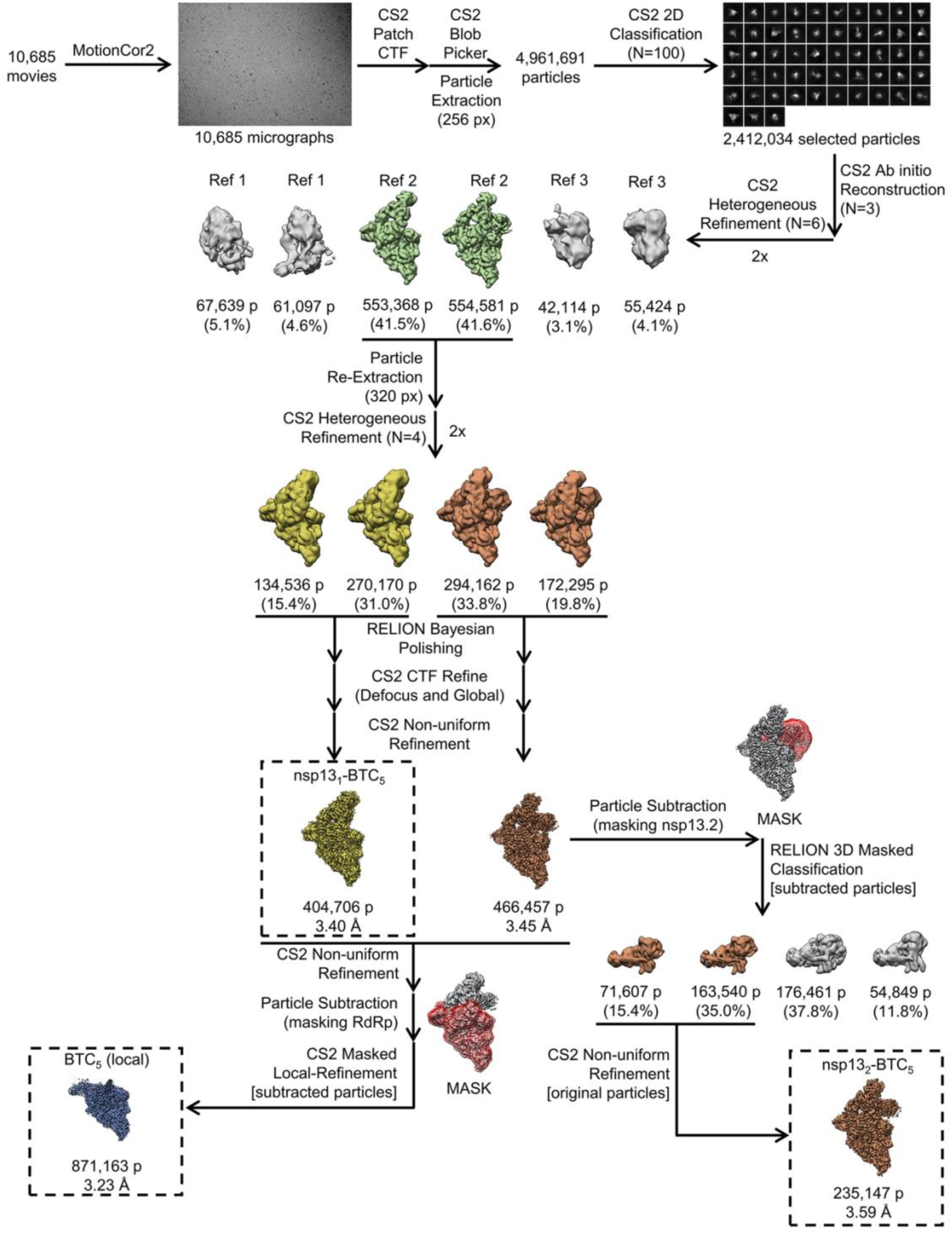
Cryo-EM processing pipeline.

**Fig. S3.**
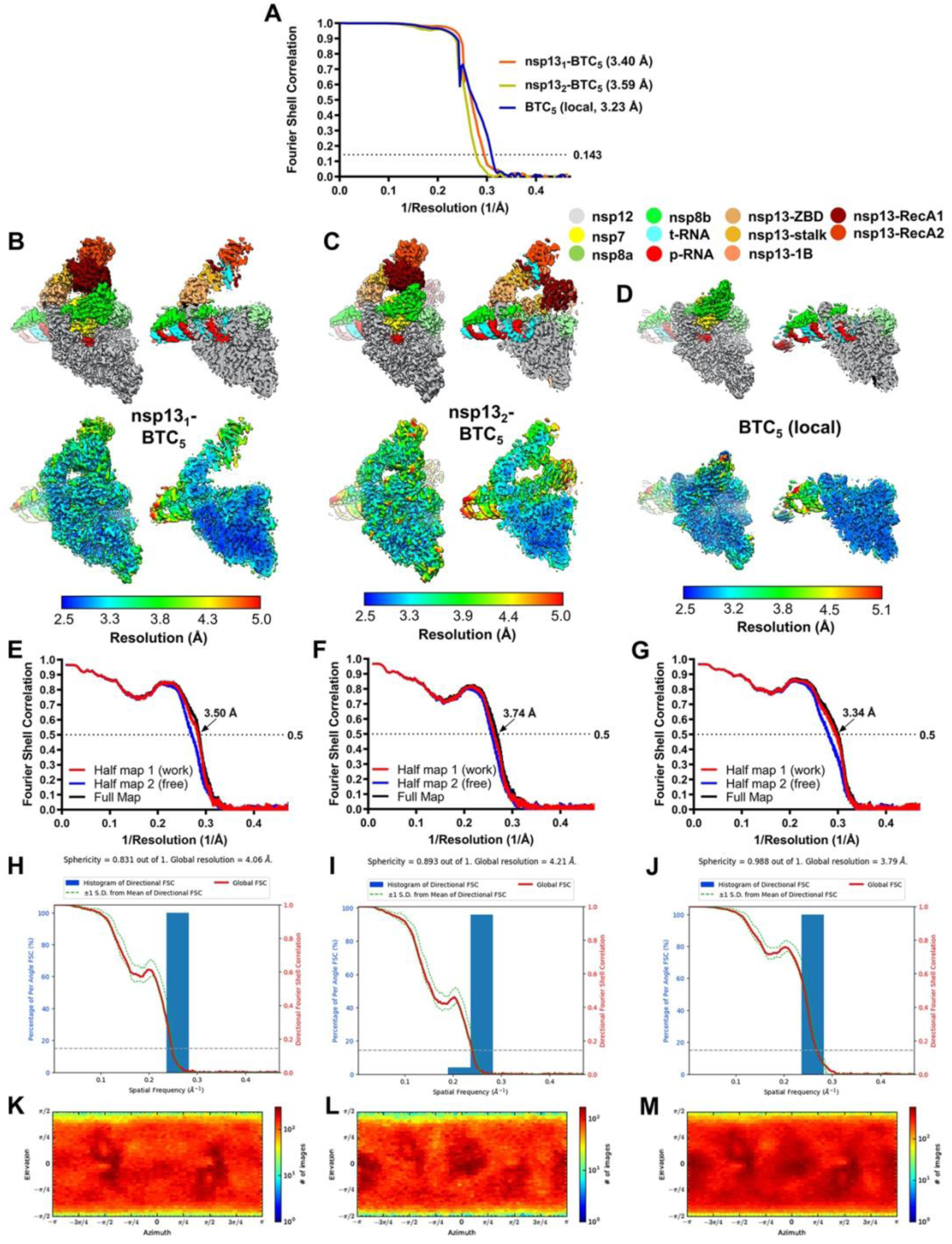
Cryo-EM analysis. **A.** Gold-standard FSC plots for nsp13_1_-BTC_5_, nsp13_2_-BTC_5_, and BTC_5_(local), calculated by comparing two independently determined half-maps from cryoSPARC (8). The dotted line represents the 0.143 FSC cutoff. **B-D.** Cryo-EM reconstructions filtered by local resolution(11). The view on the right is a cross-section. *(top)* Colored by subunit according to the color key. *(bottom)* Color by local resolution (key on the bottom). **B.** Nsp13_1_-BTC_5_. **C.** Nsp13_2_-BTC_5_. **D.** BTC_5_(local). **E – G.** FSC calculated between the refined structures and the half map used for refinement (work, red), the other half map (free, blue), and the full map (black). **E.** Nsp13_1_-BTC_5_. **F.** Nsp13_2_-BTC_5_. **G.** BTC_5_(local). **H - J,** Directional 3D Fourier shell correlation plots, calculated by 3DFSC(25). **H.** Nsp13_1_-BTC_5_. **I.** Nsp13_2_-BTC_5_. **J.** BTC_5_(local). **K – M.** Particle angular distribution plots calculated in cryoSPARC. Scale shows the number of particles assigned to a particular angular bin. Blue, a low number of particles; red, a high number of particles. **K.** Nsp13_1_-BTC_5_. **L.** Nsp13_2_-BTC_5_. **M.** BTC_5_(local).

**Fig. S4.**
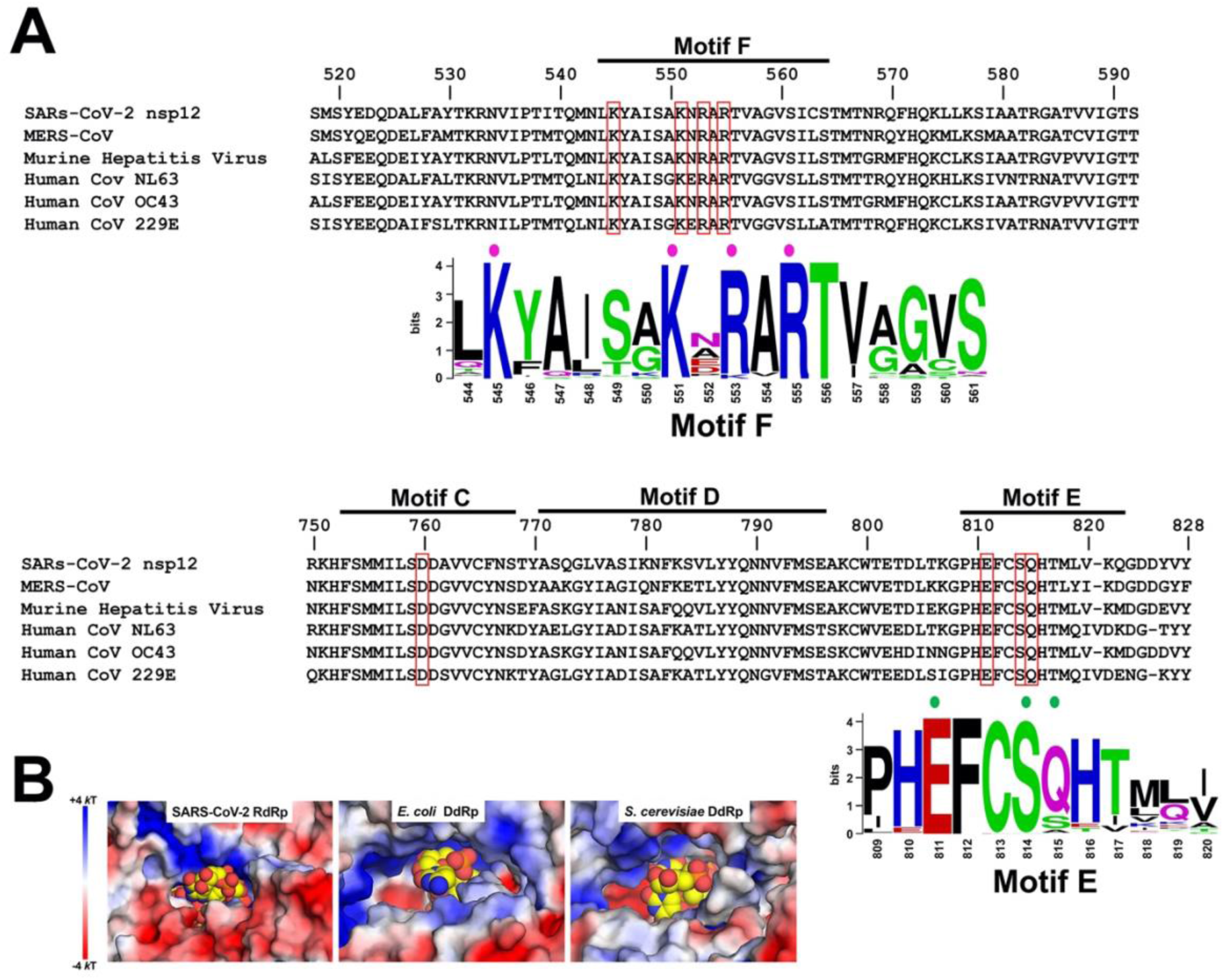
Sequence conservation of nsp12 homologs and NTP-entry tunnel environment. **A.** Sequence alignment of nsp12 homologs from six pathogenic and model CoV family members, covering RdRp motifs (27) (motifs F, C, D, and E denoted at the top of the sequence alignment) architecturally important for the NTP-entry tunnel. Selected residues discussed in the text are highlighted (red outlines). Sequence logos(28) for motif F and motif E are shown, with residues that interact with the backtracked RNA highlighted (colored dots above; see Figure 4). The sequence logos were generated from an alignment of 97 RdRp sequences from α-, β-, γ-, and δ-CoVs (Data S1). **B.** Views from the outside into the NTP-entry tunnels of the SARS-CoV-2 BTC (*left*), an *E. coli* DdRp BTC [PDB ID: 6RI9; (29)] and an *S. cerevisiae* DdRp BTC [PDB ID: 3GTP; (30)]. Protein surfaces are colored by the electrostatic surface potential [calculated using APBS; (31)]. Backtracked RNA is shown as atomic spheres with yellow carbon atoms.

**Fig. S5.**
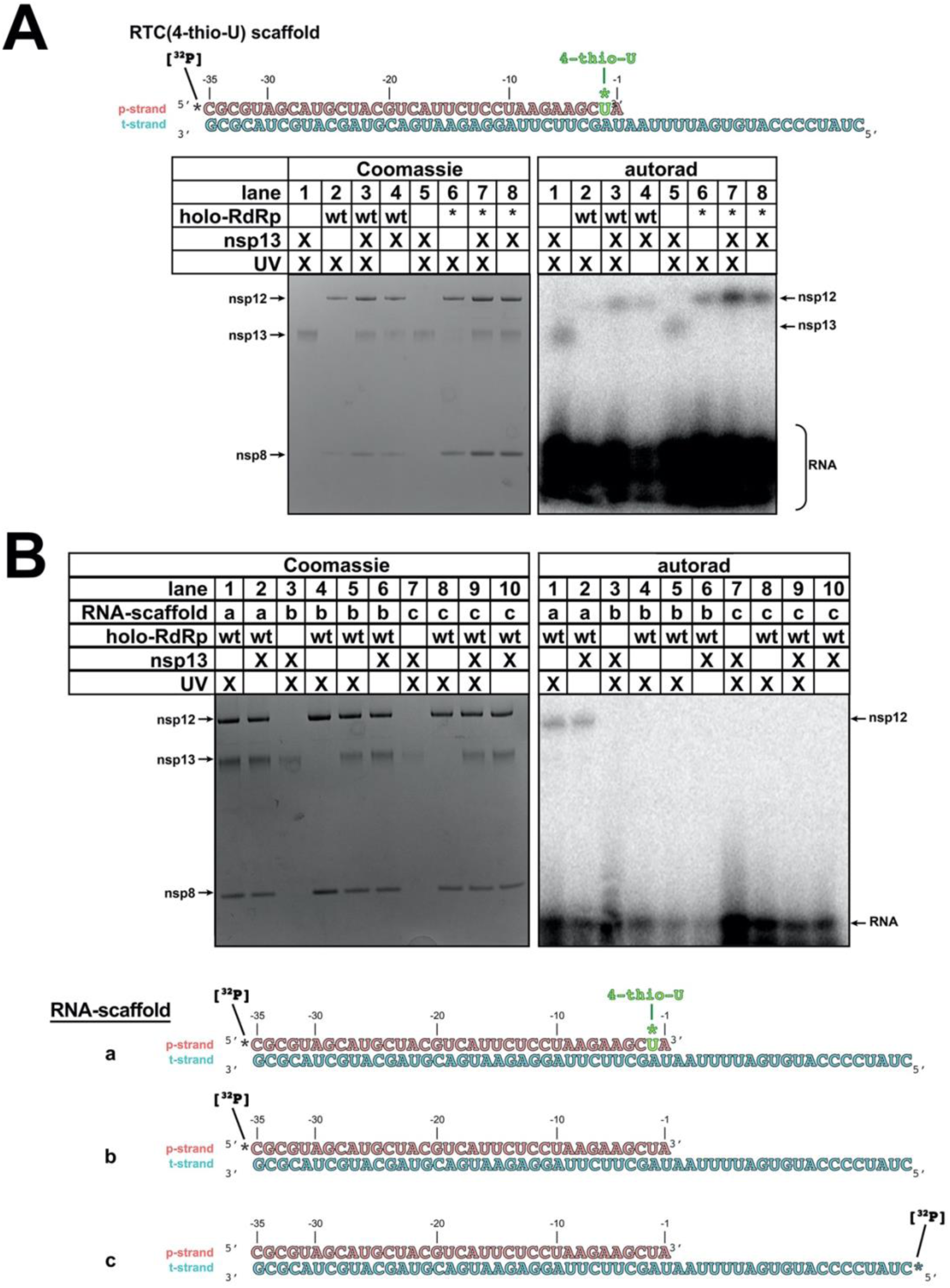
4-thio-U crosslinking analysis. **B.** Protein-RNA crosslinking analysis: The 5’-[^32^P]-labelled RTC(4-thio-U)-scaffold and the indicated proteins were incubated along with 2 mM ATP (present in every lane), exposed to UV as indicated, then analyzed by SDS polyacrylamide gel electrophoresis and autoradiography. The positions of nsp8, nsp12, and nsp13 bands are indicated. Lanes 1 and 5, containing nsp13 only, are identical controls indicating uniform UV exposure across the samples. Holo-RdRp(*) denotes the nsp12-D760A substitution that facilitates backtracking (see Figure S1A). The two panels show the same SDS polyacrylamide gel (left panel, Coomassie stained; right panel, visualized by autoradiography). **C.** Protein-RNA crosslinks are specific. Lanes 1, 2; Analysis using the 5’-[^32^P]-RTC(4- thio-U)-scaffold (RNA-scaffold ’a’ shown on the bottom). Crosslinking to nsp12 serves as a positive control for the crosslinking reaction. Lanes 3-6; Analysis using RNA-scaffold ’b’ (RTC-scaffold with 5’-[^32^P]-labelled p-RNA). Lanes 7-10: Analysis using RNA-scaffold ’c’ (RTC-scaffold with 5’-[^32^P]-labelled t-RNA). The complete absence of protein-RNA crosslinks in lanes 3-10 indicates that the observed protein-RNA crosslinks arise from the 4-thio-U site-specifically incorporated in the p-RNA of the RTC(4-thio-U)- scaffold.

**Fig. S6.**
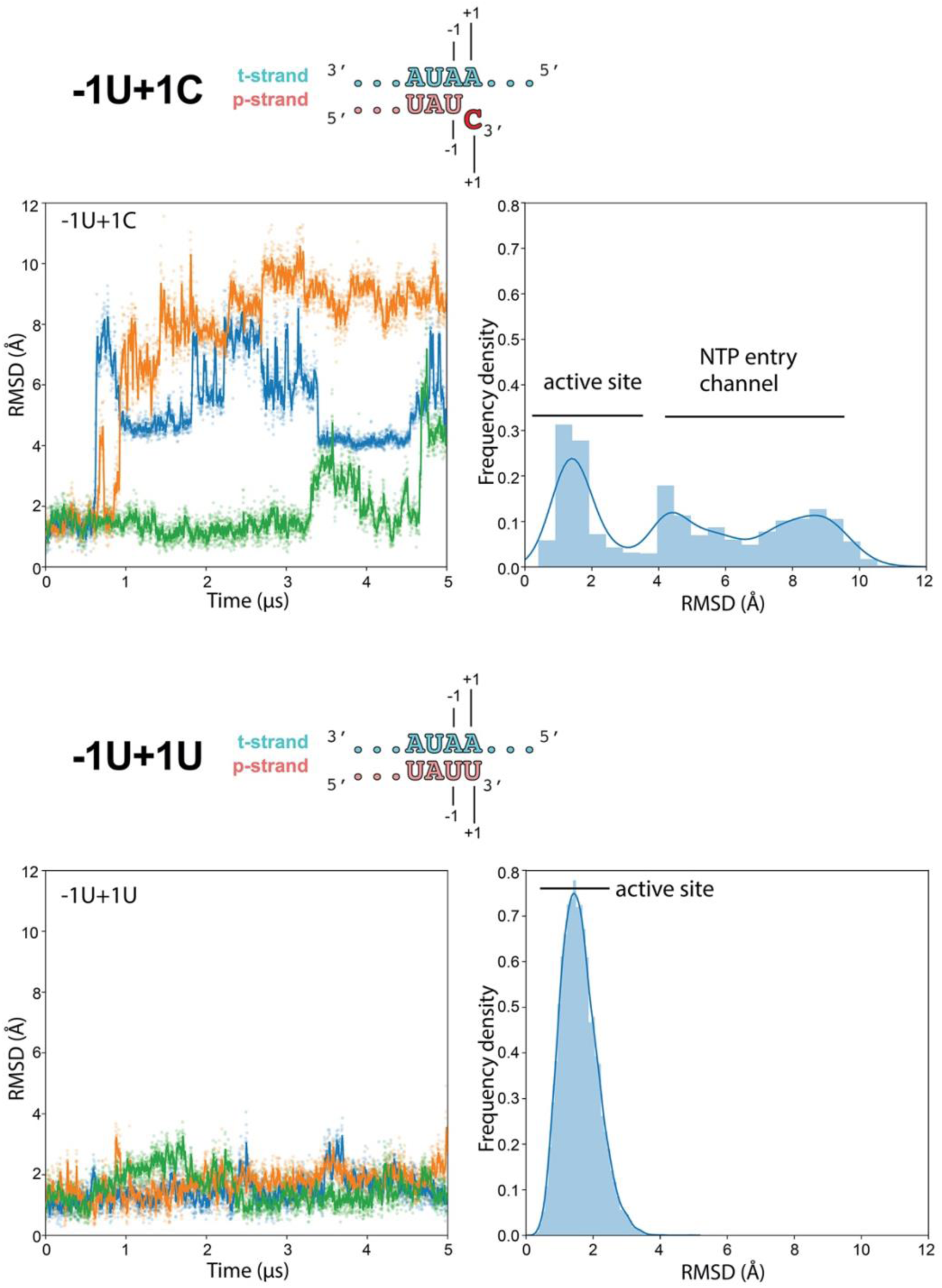
Molecular dynamics simulations of nsp13_2_-BTC_1U+1C_ vs. nsp13_2_-BTC_-1U+1U_. Molecular dynamics simulations of the nsp13_2_-BTC_−1U+1C_ (*top*) and nsp13_2_-BTC_-1U+1U_ (*bottom*) complexes. The schematics illustrate the active-site proximal nucleotides in each modeled complex. Each complex was simulated with 3 replicates. RMSD values plotted as a function of time represent the heavy-atom RMSD of the +1 nucleotide of the p-RNA (+1C for nsp13_2_-BTC_-1U+1C_ or matched +1U for nsp13_2_-BTC_-1U+1U_) compared with the starting configuration (see Methods). The RMSD histograms (plotted on the right) are aggregates of all 3 replicates. (*top*) Nsp13_2_-BTC_-1U+1C_. As shown in Figure 5C, the mismatched p-RNA +1C spends about 60% of the time frayed from the t-RNA +1A and near or in the NTP-entry tunnel (RMSD ≥ ∼3.5 Å). (*bottom*) Nsp13_2_-BTC_-1U+1U_. With the p-RNA +1U matched with the t-RNA +1A for Watson-Crick base pairing, the p-RNA +1U does not fray and spends all of its time in the vicinity of the RdRp active site and base paired with the t-RNA.

## SUPPLEMENTAL DATA FILES

**Data File S1.** Sequence alignment (Clustal format) of α- and β-CoV nsp12 sequences.

